# Dynamic Localization of Leukemia Stem Cells via CXCL12 Regulates Leukemia Progression

**DOI:** 10.1101/2024.06.29.601323

**Authors:** Chen Wang, Yi Pan, Ruochen Dong, WenXuan Zhou, XiaDuo Meng, Yao Shi, Ravi Nistala, Richard D. Hammer, Linheng Li, XunLei Kang

## Abstract

The processes that govern leukemia progression and remission are poorly understood. Our research reveals that the CXCL12 gradient, traditionally associated with LSC quiescence and survival, critically determines LSC localization and associated behavior. Specifically, CXCL12 guides LSCs to either the quiescent niche in the metaphysis, characterized by N-cadherin-expressing mesenchymal stromal cells (N-cad^+^ MSCs), or the proliferative niche in the central marrow (CM), marked by sinusoidal endothelial cells and associated stromal cells. We identified that the CXCL12 gradient is finely regulated by the interplay between dipeptidyl peptidase 4 (DPP4) on LSCs and glypican-3 (GPC3) on N-cad^+^ MSCs. DPP4 deactivates CXCL12, while GPC3 inhibits DPP4, resulting in a higher CXCL12 concentration in the metaphysis and a lower concentration in the CM. This differential gradient facilitates leukemia progression by promoting LSC quiescence and survival in the metaphysis versus relative proliferation and apoptosis in the CM. Depletion of *Dpp4* from LSCs or *Cxcl12* from N-cad^+^ MSCs disrupts this gradient, mobilizing LSCs from the metaphysis to the CM and significantly hindering leukemia development. Our findings redefine the role of CXCL12 in LSC behavior and provide a clearer understanding of leukemia progression. This novel insight highlights the potential for targeted therapeutic strategies that disrupt the CXCL12 gradient to treat minimal residual LSCs, offering a promising path toward a lasting cure for acute myeloid leukemia (AML).

## Introduction

Acute myeloid leukemia (AML) is a prevalent form of acute leukemia in adults and is characterized by the presence of leukemic stem cells (LSCs) ^1^. LSC are responsible for the initiation, progression, and relapse of leukemia. One key aspect of AML pathogenesis is the trafficking and localization of LSCs to the bone marrow (BM), where they interact with the BM niche ^2,3^. Understanding the mechanisms underlying LSC trafficking and localization in the BM can provide valuable insights into AML pathobiology and aid in the development of targeted therapies ^4,5^.

Minimal residual LSCs, marked by traits such as quiescence and drug efflux potential, significantly contribute to relapse post-remission in AML, establishing them as a well-established risk factor ^6,7^. Thus, identifying their niche emerges as a promising avenue towards achieving a permanent cure for AML. The BM niche consists of various components, including extracellular matrix, stromal cells, and cytokines, which collectively regulate LSC behavior ^8–11^. Among these factors, C-X-C motif chemokine ligand 12 (Cxcl12) and its receptor CXCR4 are of particular significance due to their suggested role in leukemia cell proliferation and migration ^12–14^. Disrupting the Cxcl12-CXCR4 signaling axis enhances chemotherapy efficacy and reduces leukemic cell adhesion and migration within the BM microenvironment ^12,15,16^. However, the precise role of Cxcl12 and its origin from BM niche cells during leukemogenesis is complex and debated, warranting further investigation^15,17–19^.

Dipeptidyl peptidase 4 (DPP4) is an important regulator of CXCL12 and has been implicated in modulating hematopoietic stem cell (HSC) engraftment ^20^. DPP4 cleaves -Ala or -Pro dipeptides off the N-terminus of cytokines/chemokines, including Cxcl12, thereby modifying their biological activities^20,21^. Recent studies have identified DPP4 as a potential therapeutic target in leukemia treatment, highlighting the need to elucidate its role in LSC trafficking and localization within the BM microenvironment ^22–25^.

Glypican-3 (GPC3), a member of the glypican family, is known to regulate cell proliferation and survival during development ^26–28^. It is a natural inhibitor of DPP4 and maintains hematopoietic stem and progenitor cells (HSPCs) within the BM ^29,30^. However, the impact of GPC3-mediated inhibition of DPP4 on LSC activity and localization remains unknown. In this study, we identified two key LSC niche areas and highlighted the importance of N-cadherin-expressing mesenchymal stromal/stem cells (N-cad^+^ MSCs) in the endosteal region of the trabecular bone^31^ as crucial contributors to LSC maintenance. Additionally, we explored the significant role of the DPP4-GPC3 interplay in regulating LSC localization and properties through controlling the CXCL12-gradient across different niches in the BM.

## Results

### *Dpp4* deficiency confines AML cells to the BM

Our group recently identified DPP4 as a potential therapeutic target for AML treatment ^22^. Strikingly, our experiments revealed that *Dpp4* deletion alters the distribution of AML cells in the BM in both MLL-AF9 and AML-ETO9a models. AML cells (*Dpp4^+/+^*) predominantly accumulate in the metaphysis area, particularly the proximal metaphysis (PM), within 3 to 5 weeks following transplantation, when the AML cell percentage in the BM is below 50%. However, *Dpp4* deletion results in a more even distribution of AML cells across the BM, with relative enrichment in the central marrow (CM) (**Fig 1a-c; Extend Data Fig 1**). Given the dramatic decrease of peripheral AML cells after *Dpp4* deletion^22^, we hypothesize that *Dpp4* deletion ‘confines’ AML cells in the BM.

**Figure 1.**
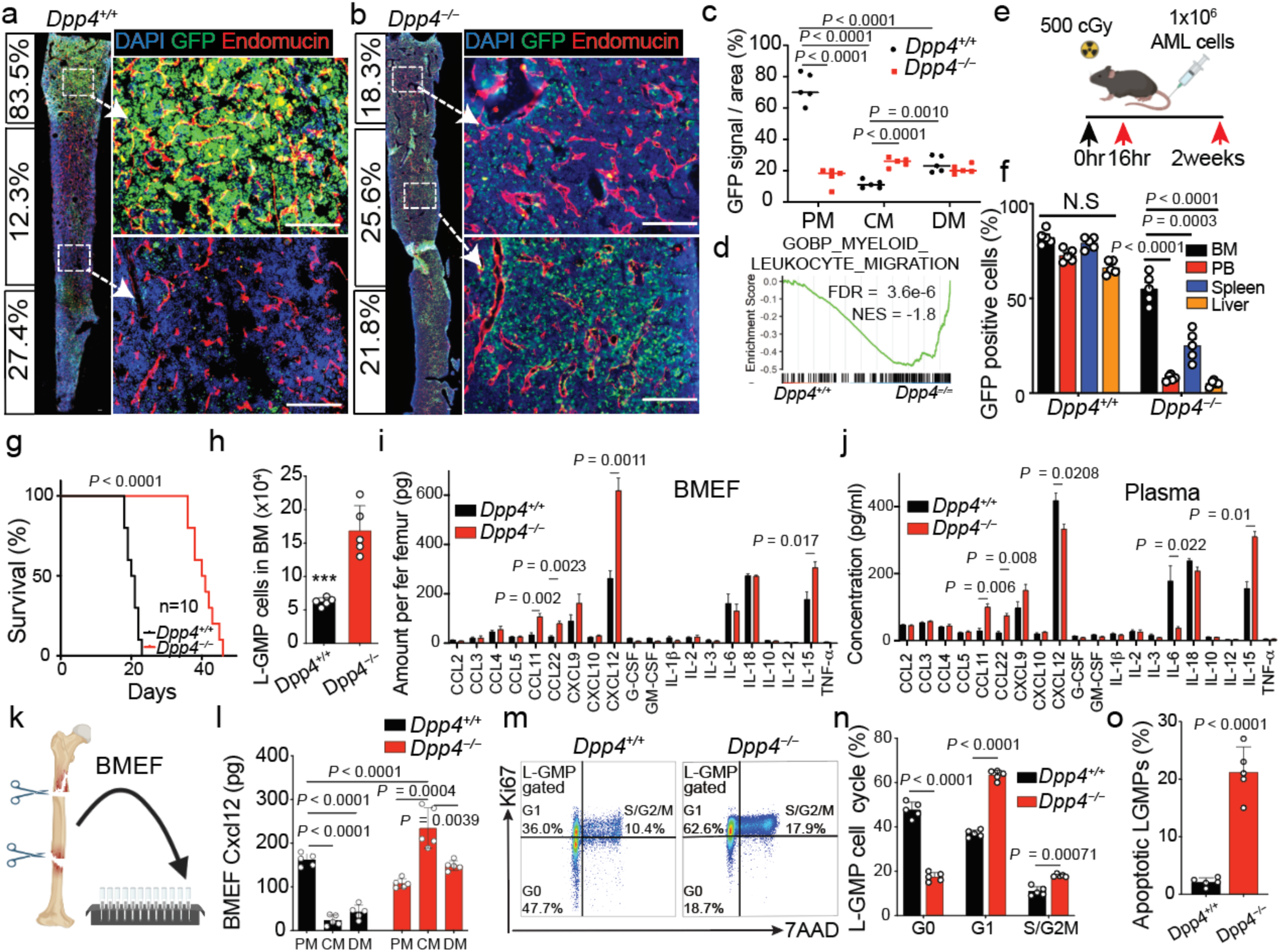
*Dpp4* deficiency alters CXCL12 gradient and regulates LSC location and properties. (a-b) Representative images of BM sections from *Dpp4*^+/+^ (a) or *Dpp4*^−/−^ (b) AML-SCs transplanted mice. Green, GFP^+^ AML cells; red, endomucin staining blood vessels; blue, DAPI for nuclei. The calculated proportion of GFP^+^ signals in each of the three anatomical BM areas is listed to the left of the images. The dashed box indicates the area of focus. Scale bars, 100μm. (c) Summary of GFP^+^ signals in each BM area (PM, CM, DM), showing that *Dpp4*^+/+^ AML cells are more heavily concentrated in the PM, while *Dpp4*^−/−^ AML cells are more evenly distributed throughout all BM areas. n = 5, BM sections from 5 mice. (d) GSEA showing negative enrichment for myeloid leukocyte migration in *Dpp4*^−/−^ AMLs, NES, normalized enrichment score; FDR, false discovery rate. (e) Schematic representation of the MLL-AF9 transplantation model for homing and engraftment assays. 1x10^6^ primary AML cells were transplanted into sub-lethally irradiated recipients, and the proportions of GFP^+^ AML cells were evaluated at the indicated time points, respectively. (f) Engraftment efficiency measured by comparison of the proportions of GFP^+^ *Dpp4*^+/+^ or *Dpp4*^−/−^AML cells in BM, PB, spleen, and liver 2 weeks post transplantation. (g) Survival curve of secondary transplanted mice receiving 1 million *Dpp4*^+/+^ or *Dpp4*^−/−^ MLL-AF9 AML cells (n = 10 mice; P< 0.001, log-rank test). (h) LSC population (IL-7R^−^Lin^−^Sca-1^−^c-Kit^+^CD34^+^FcψRII/III^+^, GMP-like leukemic cells, L-GMP) measured by flow cytometry in the BM of *Dpp4*^−/−^ and *Dpp4*^+/+^ AML mice (n = 5 mice). (i-j) Cytokine protein levels were measured in BMEF (i) and plasma (j) of *Dpp4*^+/+^ and *Dpp4*^−/−^ AML cells transplanted mice using the LEGEND plex Muti-Analyte Flow Assay Kit. (n = 3 mouse samples). (k-l) Cartoon of BM structure anatomical partition (k) and summary (l) of Cxcl12 levels were measured in each BM area (PM, CM, and DM) from *Dpp4*^−/−^ and *Dpp4*^+/+^ AML mice. (m-n) Cell cycle analysis of MLL-AF9 AML-SCs in *Dpp4*^+/+^ and *Dpp4*^−/−^ mice (n = 5 mice per group) by flow cytometry. (o) Comparison of apoptotic LSCs in the BM between *Dpp4*^+/+^ and *Dpp4*^−/−^ MLL-AF9 mice (n = 5 mice). Error bars, s.e.m. Data were analyzed using a two-tailed Student’s t-test for pairwise analysis. When comparing 3 or more groups, one-way or two-way ANOVA was used with Fisher’s test for post hoc analyses.

To validate this hypothesis, we performed RNA-seq analysis on AML cells from MLL-AF9 mouse model, finding that DPP4 knockout (*Dpp4^−/−^*) AML cells showed reduced gene signatures associated with myeloid leukocyte migration (**Fig 1d and Extend Data Fig 2a and b**). We then transplanted one million *Dpp4^+/+^*or *Dpp4^−/−^* AML cells into wild-type recipient mice (**Fig 1e**). Although the initial homing (16 hours post-transplantation) of *Dpp4^−/−^* AML cells was comparable to *Dpp4^+/+^* cells, engraftment analysis (2 weeks post-transplantation) showed that *Dpp4^−/−^* AML cells were predominantly localized in the BM, while *Dpp4^+/+^* cells were more evenly distributed across different hematopoietic organs (**Fig 1f and Extend Data Fig 2c**).

Notably, mice transplanted with *Dpp4^−/−^* MLL-AF9 AML cells survived twice as long as those with *Dpp4^+/+^* AML cells (40.6 days vs. 20.5 days, **Fig 1g and Extend Data Fig 2d and e**). Furthermore, the BM of *Dpp4^−/−^* AML cells transplanted mice (referred to as *Dpp4^−/−^* AML mice) contained significantly more LSCs, specifically GMP-like leukemic cells (L-GMP, IL-7R^−^ Lin^−^ Sca-1^−^ c-Kit^+^ CD34^+^ FcγRII/III^+^), compared to *Dpp4^+/+^* AML mice (a 2.71-fold increase, **Fig 1h and Extend Data Fig 2f**) ^32^. Using AML cells transformed by the AML1-ETO9a mutation ^33^, similar results were observed: *Dpp4^−/−^* AML1-ETO9a mice exhibited significantly longer survival (73.5 days vs. 40.5 days, **Extend Data Fig 2g**), with more restricted BM localization and a higher proportion of LSC progenitor cells (**Extend Data Fig 2h-j**), compared to those transplanted with *Dpp4^+/+^* AML1-ETO9a AML cells. These findings strongly suggest *Dpp4* deletion confines AML cells to the BM, highlighting DPP4’s crucial role in regulating LSC trafficking and disease progression.

### Reversed Cxcl12 gradient in *Dpp4^−/−^* AML mice alters AML cell trafficking and LSC properties

To understand the altered AML cell trafficking due to *Dpp4* deletion, we investigated potential physical barriers between AML cells and blood vessels. Immunofluorescence analysis showed that both GFP-labeled *Dpp4^+/+^* and *Dpp4^−/−^* AML cells were equidistant from endomucin-labeled vessels throughout the BM, indicating unimpeded vascular access (**Fig 1a, b; Extend Data Fig 1a, b; Extend Data Fig 3**) ^34,35^. We then examined cytokine and chemokine levels, focusing on DPP4 substrates and factors influencing AML cell trafficking ^36–40^. Significant differences were found in the concentrations of CCL11, CCL22, CXCL12, IL-6, and IL-15 between *Dpp4^+/+^*and *Dpp4^−/−^* mice. Notably, CXCL12 exhibited a reversed gradient from plasma to BM in *Dpp4^−/−^* mice compared to *Dpp4^+/+^* mice (**Fig 1i and j; Extend Data Fig 4a-c**). This reversal was specific to CXCL12, as other cytokines and chemokines showed no such gradient changes. Additionally, flow cytometry revealed similar CXCR4 expression on both *Dpp4^+/+^* and *Dpp4^−/−^* LSCs (**Extend Data Fig 4d**), indicating that these effects were mediated specifically by CXCL12.

To assess the functional consequences of the altered CXCL12 gradient regulated by DPP4, we conducted chemotaxis assays (**Extend Data Fig 4e top panel**). In the absence of CXCL12, there was no significant difference in migration between *Dpp4^+/+^*and *Dpp4^−/−^* AML cells **(Extend Data Fig 4e (i))**. However, in the presence of CXCL12, *Dpp4^−/−^* AML cells exhibited significantly reduced migration (**Extend Data Fig 4e (ii-iii), f**), suggesting that AML cells are attracted to and confined within areas of high CXCL12 concentration. To further validate DPP4’s role in AML cell trafficking via CXCL12 regulation, we administered exogenous CXCL12 intravenously to both *Dpp4^+/+^*and *Dpp4^−/−^* AML mice. The administration of CXCL12 resulted in a substantial increase in peripheral AML cells in *Dpp4^+/+^* AML mice compared to *Dpp4^−/−^*AML mice **(Extend Data Fig 4g, h)**.

Examining CXCL12 concentration in different BM regions (PM, CM, and DM), we found that CXCL12 predominantly accumulated in the PM area of *Dpp4^+/+^* AML mice, whereas *Dpp4^−/−^*AML mice showed higher CXCL12 levels in the CM area, with less pronounced regional differences (**Fig. 1k, l**). These findings suggest that CXCL12 is a key messenger in AML cell trafficking, with DPP4 deficiency reversing the CXCL12 gradient, thereby altering LSC trafficking and localization.

To evaluate the functional impact of altered LSC localization, we compared the cell cycle profile and apoptosis of LSC in the BM of *Dpp4^+/+^*and *Dpp4^−/−^* AML mice. The percentage of L-GMP cells in the G0 phase was reduced by 63.4%, whereas the G1 phase showed a 1.7-fold increase in *Dpp4^−/−^* AML mice compared to that in *Dpp4^+/+^*AML mice (**Fig. 1m, n**). This indicates a substantial decrease in LSC quiescence in *Dpp4^−/−^* AML mice, consistent with the previously observed reduction in stemness of *Dpp4^−/−^*AML cells ^22^. Additionally, there was a significantly higher apoptosis rate in LSCs within the BM of *Dpp4^−/−^*mice compared to *Dpp4^+/+^* AML mice (**Fig. 1o, Extend Data Fig 4i**), aligning with the established role of DPP4 in AML cell survival^22,25^. These findings highlight the pivotal role of DPP4 in determining AML cell location and distribution within the BM by altering the CXCL12 gradient. This suggests that LSC properties are influenced by their location regulated through CXCL12, rather than by direct action of CXCL12.

### N-cad^+^ MSCs support LSCs within the BM through CXCL12

To understand which niche cells and factors contribute to LSCs accumulation in the PM area of the BM, we performed a single-cell RNA sequencing (scRNA-seq) on non-hematopoietic BM cells from *Dpp4^+/+^* AML mice three weeks post-transplantation. Single-cell suspensions were prepared from BM samples using a combination of grinding and collagenase-dispase treatment, and viable non-hematopoietic cells (CD45^−^GFP^−^) were sorted for scRNA-seq analysis (**Fig. 2a**) ^41,42^. Through the analysis of 3764 cells (median of 16053.5 molecules and 3556.5 genes per cell), we identified 18 distinct clusters (0−17), spanning mesenchymal stem/stromal cells (MSC), osteolineage cells (OLC), chondrocytes (Chondro), fibroblasts (Fibro), endothelial cells (EC), pericytes, and possible transitional states (**Fig. 2b, c**) ^43,44^.

**Figure 2.**
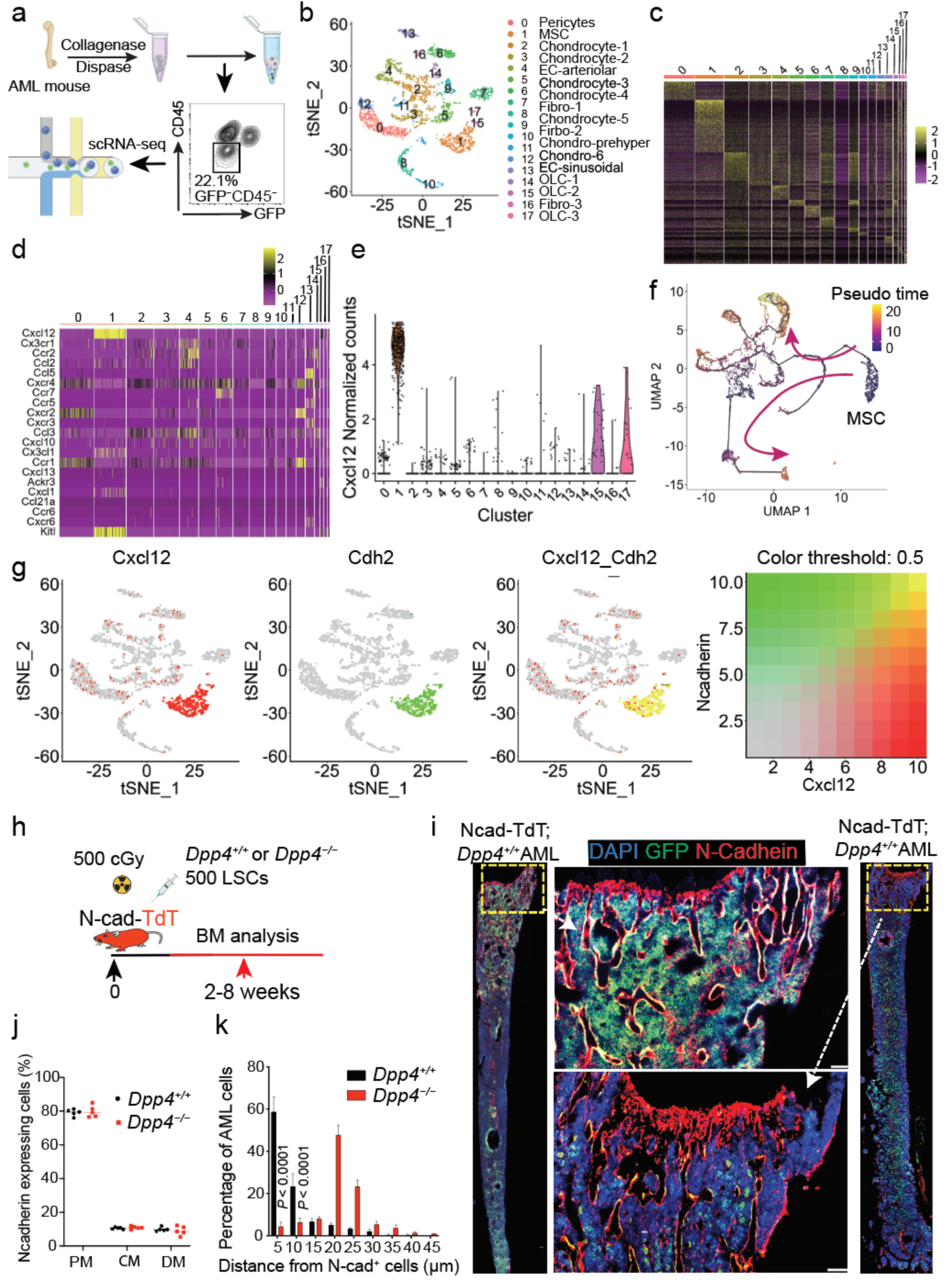
N-cad^+^ BMSCs support LSCs within the BM through *Cxcl12*. (a) Schematic diagram for single cell sequencing. Single-cell library was prepared from BM CD45^−^GFP^−^ cells and sent for scRNA-seq analysis. (b-c) Eighteen bone marrow stromal cell clusters. t-Distributed stochastic neighbor embedding (t-SNE) of 3764 bone marrow niche cells, annotated post hoc and colored by clustering (b) and expression of top differentially expressed genes (rows) across the cells (columns) in each cluster (c). (d-e) Expression of key niche factors, cytokines, and chemokines across all clusters, in which *Cxcl12* is enriched in MSC (cluster 1). (f) Trajectory analysis suggests that MSCs serve as the origin of BM niche cells. (g) Coexpression pattern of *Cxcl12* and N-Cadherin. Red, green, and yellow denote *Cxcl12* expression, N-cadherin expression, and *Cxcl12*-N-cadherin coexpression, respectively. Significant coexpression is observed in MSC (cluster 1). (h) Schematic representation of transplanting 500 LSCs into the sub-lethally irradiated N-cad-tdTomato (N-cad-TdT) mice and performing BM imaging. (i) Representative images of BM sections of *Dpp4*^+/+^ (left) and *Dpp4*^−/−^ (right) AML cell localization in the BM of N-cad-TdT mice. Green, GFP^+^ AML cells; red, N-cad^+^ BMSCs; blue, DAPI for nuclei. The dashed box indicates the area of focus. Scale bars 100μm. (j) Summary of N-cad-TdT cells in each BM area (PM, CM, and DM) showing that there is no significant difference of N-cad^+^ BMSCs throughout all BM areas between *Dpp4*^+/+^ and *Dpp4*^−/−^ AML cell transplantation. n = 5, BM sections from 5 mice. (k) Distance between AML cells and N-cad^+^ cells. The numbers on the x axis indicate intervals of 5 μm (5 indicates the interval 0–5; 10 indicates 5–10, and so on). p value by the two-sample Kolmogorov-Smirnov (KS) test. Error bars, s.e.m. Data were analyzed using a two-tailed Student’s t-test for pairwise analysis. When comparing 3 or more groups, a one-way or two-way ANOVA was used with Fisher’s test for post hoc analyses.

Specifically, we identified one pericyte cluster, one MSC cluster, six Chondro clusters, two EC clusters, four Fibro clusters, and three OLC clusters. Cxcl12 was predominantly (94.8%) expressed by the MSC cluster (cluster 1), and MSC differentiates into other lineages of cells as shown by trajectory analysis (**Fig. 2d-f**) ^45^. Notably, another well-known niche factor for hematopoietic stem cells (HSCs), stem cell factor (SCF, also known as Kit ligand, Kitl), was also enriched in this cluster (**Fig. 2d, e and Extend Data Fig 5a**).MSCs, crucial components of the BM hematopoietic microenvironment, are heterogeneously constituted with several reported and overlapping fractions, such as LepR^+^, N-cad^+^, Prx-1^+^, and Nes^+^ MSCs ^31,45–51^. According to the reported distribution of each subgroup of MSCs in genetic lineage tracing studies in mice (LepR-tdTomato^49^, N-cad-tdTomato^31^, Prx-1-tdTomato^52^, and Nes-GFP^51^, respectively), we hypothesized that N-cad^+^ MSCs represent an important niche for LSCs, as the localization of N-cad^+^ MSCs closely resembled that of *Dpp4^+/+^* AML cells in the BM (**Fig. 1a)**. Colocalization analysis revealed that 95.3% (452 out of 474) of N-Cad^+^ cells expressed Cxcl12, and 59.5% (452 out of 760) of Cxcl12 was expressed by N-Cad^+^ cells, indicating that N-Cad^+^ MSCs are important sources of Cxcl12 in the AML BM niche (**Fig. 2g**). While N-Cad^+^ MSCs have been previously identified as niche cells for HSCs in the endosteum of the trabecular bone area (TBA) ^31,47^, their role in AML development remains unknown.

To further investigate the association between N-cad^+^ MSCs and LSCs, we transplanted LSCs into N-cad-tdTomato (N-cad-TdT) mice, where Tomato^+^ cells report N-cadherin expression ^31^ (**Fig. 2h**). Immunostaining confirmed that around 80% of N-cad^+^ MSCs were localized in the PM area, with some presence in the CM and DM areas of the BM. This distribution was similar between *Dpp4^+/+^* and *Dpp4^−/−^* LSCs transplanted mice (**Fig. 2i, j**). Furthermore, quantification data revealed a distinct spatial relationship: 58.6% ± 7.3% of *Dpp4^+/+^* AML cells were found within a 5μm proximity to N-cad^+^ cells, whereas only 10.5% ± 2.1% of *Dpp4^−/−^* AML cells were located within the same 5μm range from N-cad^+^ cells, manifesting a distinctive “not close to me” pattern (**Fig. 2i, k**). These data strongly suggest that N-cad^+^ MSCs in the PM area of the BM are critical niche cells for LSCs and support LSCs through CXCL12.

### The deletion of *Cxcl12* from N-cad^+^ MSCs significantly inhibits AML development

To investigate the role of CXCL12 from N-Cad^+^ cells in AML development, we transplanted LSCs into N-cad^CreER^; Cxcl12^fl/fl^ mice. Conditional knockout of *Cxcl12* from N-Cad^+^ MSCs (N-cad; *Cxcl12^−/−^*) significantly inhibited AML development, evident by the prolonged survival and reduced infiltration of AML cells in hematopoietic organs, compared to control mice (N-cad; *Cxcl12^+/+^*) (**Fig. 3a-d**). AML cells in N-cad; *Cxcl12^−/−^* mice were primarily confined to the BM (**Fig. 3e**), with an increased number of LSCs compared to control mice (1.78-fold increase) (**Fig. 3f)**. Immunostaining revealed that LSCs in N-cad; *Cxcl12^+/+^* mice predominantly resided in the PM area, while those in N-cad; *Cxcl12^−/−^*mice were evenly distributed, with higher ratios in the CM area (**Fig. 3g-i**). Similar experiments using N-cad^CreER^; Scf ^fl/fl^ mice as recipient mice did not show alterations in AML development upon SCF deficiency in N-cad^+^ MSCs **(Extend Data Fig 5b, c**), suggesting the unique role of Cxcl2 in regulation of AML development. These findings underscore the critical role of CXCL12 from N-cad^+^ MSCs in regulating AML development.

**Figure 3.**
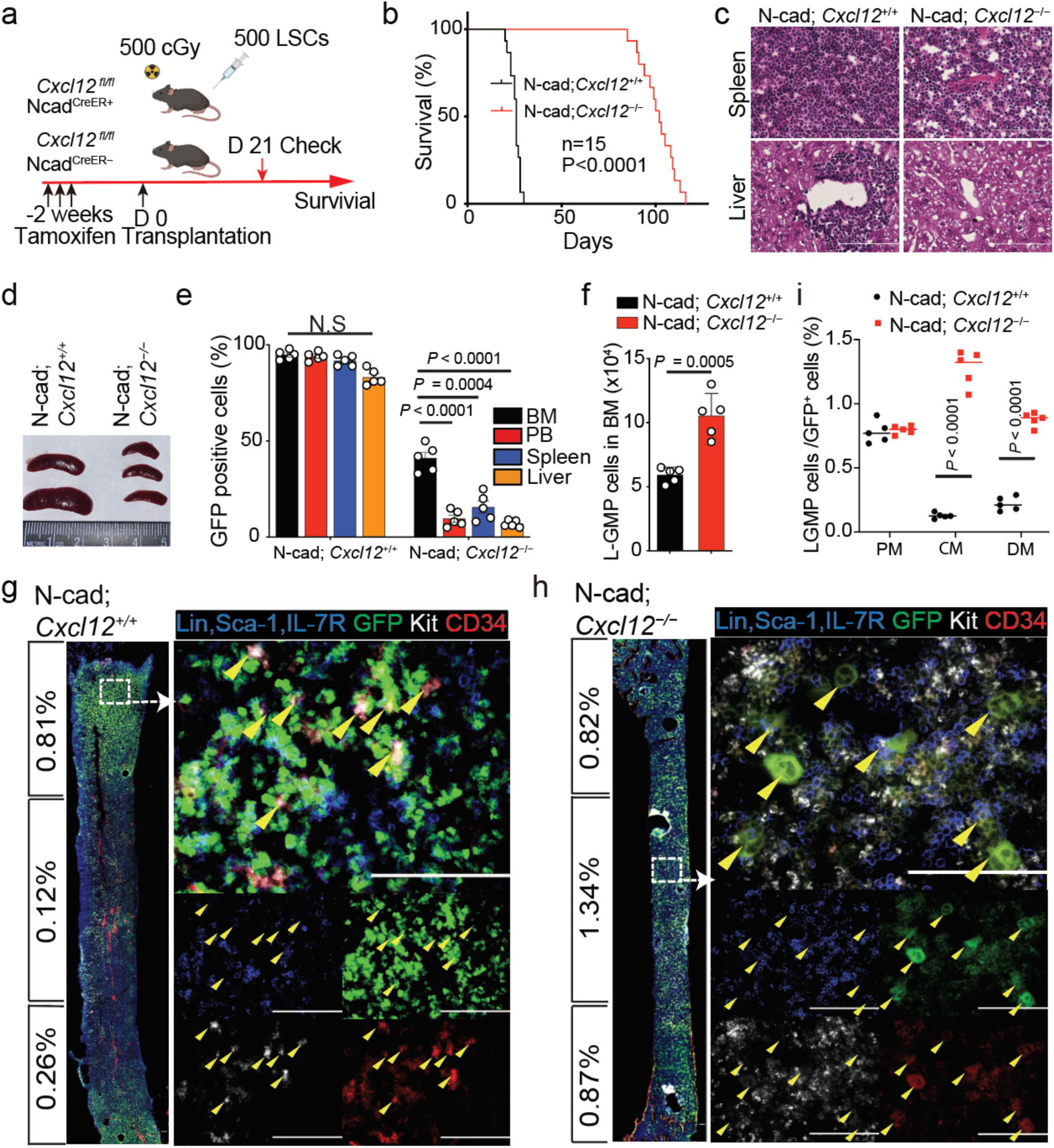
The deletion of *Cxcl12* from N-cad^+^ MSCs significantly blocks AML development. (a) Schematic representation of *Cxcl12* conditional knockout in the N-cad^+^ cell mouse model. 500 primary LSCs were transplanted into lethally irradiated recipients 14 days post tamoxifen injection. (b) Survival curve of LSC transplanted recipient N-cad; *Cxcl12*^+/+^ or N-cad; *Cxcl12*^−/−^ mice (n= 15 mice; P< 0.001, log-rank test). (c) H&E staining for spleen and liver at 6-week post transplantation. (d) Comparison of the size of spleens of the recipient N-cad; *Cxcl12*^+/+^ and N-cad; *Cxcl12*^−/−^ mice transplanted with LSCs at 6-weeks post transplantation. (e) Comparison of the proportions of GFP^+^ N-cad; *Cxcl12*^+/+^ or N-cad; *Cxcl12*^−/−^ AML cells in BM, PB, spleen, and liver 6 weeks post transplantation (n=5 mice). (f) AML-SC (L-GMP) measured by flow cytometry in the BM of N-cad; *Cxcl12*^+/+^ or N-cad; *Cxcl12*^−/−^ AML mice (n = 5 mice). (g-h) Immunostaining of L-GMP markers in the BM section of N-cad; *Cxcl12*^+/+^ (g) and N-cad; *Cxcl12*^−/−^ (h) AML mice. The calculated proportion of L-GMP cells / GFP^+^ cells in each of the three anatomical BM areas is listed to the left of the images. The dashed box indicates the area of focus. Arrow heads point to the L-GMP cells, scale Bars, 100μm. (i) Summary of distribution of AML-SCs in the PM, CM and DM areas by ImageJ. Error bars, s.e.m. *** p < 0.001. Data were analyzed using a two-tailed Student’s t-test for pairwise analysis. When comparing 3 or more groups, a one-way or two-way ANOVA was used with Fisher’s test for post hoc analyses.

### N-cad; *Cxcl12^−/−^* and *Dpp4^−/−^* AML show highly consistent signaling change

RNA sequencing of AML cells from N-cad; *Cxcl12^−/−^* mice revealed upregulation of genes associated with cell cycle regulation and metabolism, and downregulation of genes linked to stemness, migration, and NF-κB signaling **(Fig. 4a, b)**. Functional assays confirmed increased cell cycle activity and apoptosis, reduced self-renewal, and migration in LSCs from N-cad; *Cxcl12^−/−^* mice (**Fig. 4c-h**). Transcriptome analysis showed reduced activity of JAK/STAT, MAPK, and NF-κB pathways in these cells, with diminished activation of key signaling molecules (**Fig. 4i, j**). These pathways, known to be activated by the Cxcl12/CXCR4 axis, are critical for survival, cell proliferation, chemotaxis, and adhesion in various tumor cells ^53–59^.

**Figure 4.**
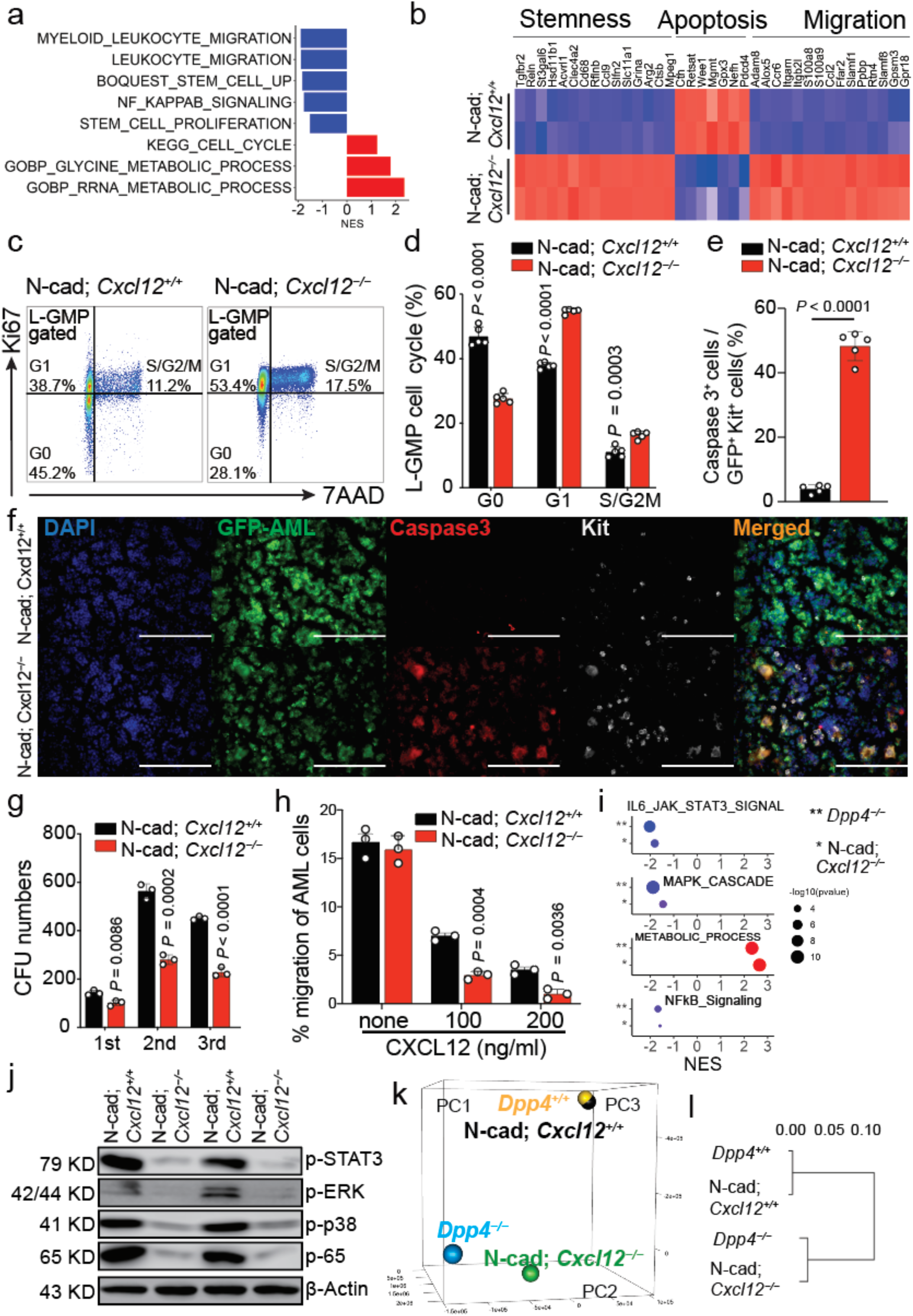
N-cad; *Cxcl12*^−/−^ and *Dpp4*^−/−^ AML mice show highly consistent signaling change. (a) Bar plot of GSEA results illustrating biological processes associated with AML cells of *Cxcl12* conditional knockout in N-Cadherin cells. It shows the significant alteration of migration, stemness, proliferation, and the cell cycle. (b) Heatmap of differentially expressed genes in N-cad; *Cxcl12*^+/+^ and N-cad; *Cxcl12*^−/−^ AML cells. (c-d) Cell cycle analysis of MLL-AF9 AML-SCs in N-cad; *Cxcl12*^+/+^ and N-cad; *Cxcl12*^−/−^ AML mice (n = 5 mice per group) by flow cytometry. (e-f) Immunofluorescent image shows the levels of cleaved caspase-3 in the BM of N-cad; *Cxcl12*^+/+^ and N-cad; *Cxcl12*^−/−^ AML mice. Green, GFP+ AML cells; red, cleaved-Caspase3, bule, DAPI; white, Kit^+^ cells; scale bars, 100μm. (g) Comparison of CFU capability of N-cad; *Cxcl12*^+/+^ and N-cad; *Cxcl12*^−/−^ AML cells during serial re-plating (2,000 cells/well, n = 3 wells). (h) *In-vitro* chemotaxis assays (described in Extend Data Fig 4E), show the migration of AML cells from N-cad; *Cxcl12*^−/−^ AML mice towards *Cxcl12* induction is significantly reduced compared to the AML cells from N-cad; *Cxcl12*^+/+^ AML mice. (i) Highly consistent GSEA change between *Dpp4*^−/−^ vs. *Dpp4*^+/+^ (**) and N-cad; *Cxcl12*^−/−^ vs. N-cad; *Cxcl12*^+/+^ (*), including down-regulation of STAT3, MAPK, and NF-kB pathways and increased metabolic processes. NES, normalized enrichment score; FDR, false discovery rate. (j) Comparison of p-STAT3, p-ERK, p-P38, and p-p65 protein levels in N-cad; *Cxcl12*^+/+^ and N-cad; *Cxcl12*^−/−^ MLL-AF9 BM cells from each of 2 AML mice. (k-l) PCA analysis (k) and Pearson distance tree (l) for AML cells from N-cad; *Cxcl12*^−/−^ and *Dpp4*^−/−^ AML mice and their pairing N-cad; *Cxcl12*^+/+^ and *Dpp4*^+/+^ AML mice. (The x axis represents 1–r (Pearson’s correlation value). Error bars, s.e.m. Data were analyzed using a two-tailed Student’s t-test for pairwise analysis. When comparing 3 or more groups, a one-way or two-way ANOVA was used with Fisher’s test for post hoc analyses.

Interestingly, the altered signaling observed is highly consistent with that of *Dpp4^−/−^* AML mice (**Fig. 4i**). Pearson distance tree and principal component analysis (PCA) revealed that *Dpp4^−/−^*AML cells and AML cells from N-cad; Cxcl12^−/−^ mice had closely related transcriptome profiles, distinct from their respective controls. This supports the collaboration between N-cad^+^ MSCs and *Dpp4^+^* AML cells in regulating CXCL12 integrity and AML development (**Fig. 4k, l**). The similarity in transcriptome profiles aligns with the resembling phenotypes observed between N-cad; *Cxcl12^−/−^* and *Dpp4^−/−^* AML (**Fig. 1, 3**).

### GPC3 specifically expressed on N-cad^+^ MSCs inhibits DPP4 and attracts LSCs to the PM

We have demonstrated the critical role of CXCL12 in orchestrating the localization of LSCs in the BM, with DPP4 known to efficiently inactivate CXCL12 ^22,60^. This indicates a mechanism by which DPP4 regulates LSC localization. However, an intriguing question remains: how do *Dpp4^+/+^* LSCs achieve close proximity to N-cad^+^ niche cells actively secreting CXCL12? We hypothesized that another factor in the BM preserves the integrity of CXCL12 directly or indirectly. To test this hypothesis, a transcriptional study was performed on 10 known Cxcl2 or DPP4 regulators in both LSCs and N-cad^+^ BMSCs (**Extend Data Fig 6**) ^29,61,62^. The study revealed that Glypican-3 (GPC3) had a dramatically higher expression level than other factors, specifically in N-cad^+^ MSCs (**Fig. 5a**). GPC3 is known as an inhibitor of DPP4, suggesting its potential role in suppressing the effect of DPP4 in AML cells and preserving CXCL12 function around N-cad^+^ MSCs. Flow cytometry analysis confirmed significantly higher cell surface expression of GPC3 in N-cad^+^ cells in the BM (**Fig. 5b**). Additionally, GPC3 was found to specifically bind to DPP4-positive AML cells *in vitro* (**Fig. 5c**). Immunostaining further detected the expression and localization of GPC3 and DPP4 in the BM (**Figs. 5d-f**). Quantification showed that N-cad^+^ cells (79.8% ± 3.3%) exhibited significantly higher GPC3 expression compared to N-cad^−^ cells (10.5% ± 2.1%) in the PM area (**Fig. 5g**). AML cells also showed significantly higher DPP4 expression than non-AML cells (**Fig. 5h**), consistent with previous findings ^22^. Interestingly, 56.5% ± 8.6% of the LSCs-enriched population (GFP^+^ Kit^+^ cells) compared to only 10.6% ± 2.2% of differentiated AML cells (GFP^+^ Kit^−^ cells), were located within 5 μm from N-cad^+^ cells (**Fig. 5i**). In these N-cad+ cells and closely located LSCs, about 73.5% of GPC3 and DPP4 were overlapped (**Fig. 5j**).

**Figure 5.**
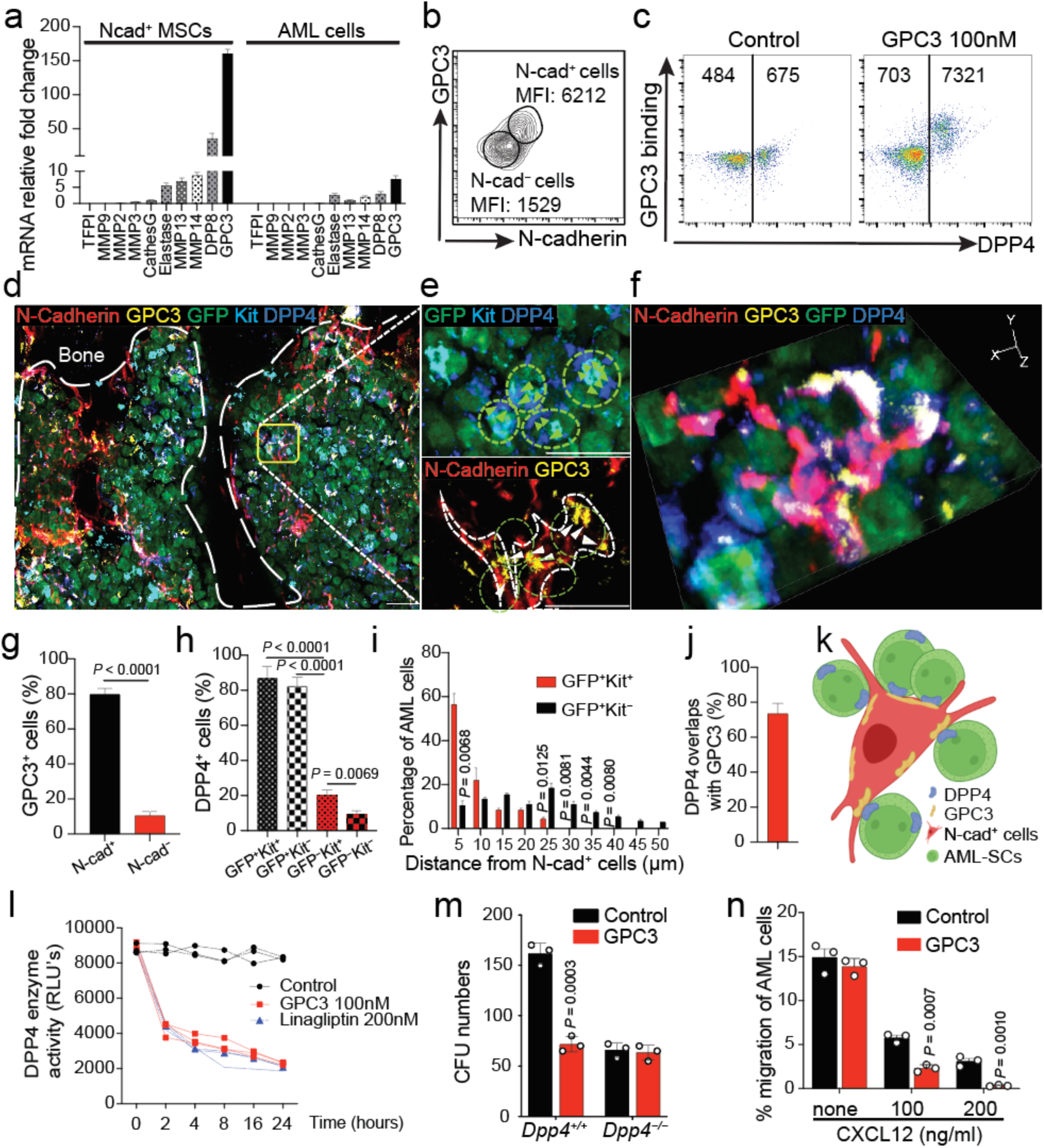
The specific expression of GPC3 on N-cad^+^ BMSC inhibits DPP4 and attracts LSCs to the PM. (a) Bar graphs compare the change in expression levels of the indicated transcripts in N-cad^+^ BMSCs and MLL-AF9 AML cells. All transcripts’ levels were normalized to levels of actin expression (n = 3 wells). (b) Contour plots show that N-cad^+^ BM cells have significantly higher cell surface expression of GPC3 than N-cad^−^ BM cells. GPC3 MFIs of N-cad^+^ and N-cad^−^ populations have been indicated. (c) Flow cytometry analysis of recombinant GPC3 (100 ng) binding to DPP4^+^ BM AML cells, MFIs are indicated. (d-e) Representative BM section images of GFP-labeled *Dpp4*^+/+^ AML cells in the N-cad-TdT AML mouse trabecular bone region. The dashed box indicates the area of focus. Dash line indicates the bone structure; scale bar, 20μM. (f) Selected 3D image of DPP4 and GPC3 interaction. Green, GFP^+^ AML cells; red, N-cad^+^ BMSCs; blue, DPP4; yellow, GPC3. (g-h) Statistic analysis of expression and localization of GPC3(g) and DPP4(h). (i) Relative distance between GFP^+^Kit^+^ AML cells and GFP^+^Kit^−^ AML cells to N-cad^+^ cells (n = 100 GFP^+^Kit^+^ AML cells, n = 80 GFP^+^Kit^−^ AML cells). n=3 mice for each distance quantification dataset. (j) Percentage of DPP4^+^ AML cells overlap with GPC3^+^ N-cad^+^ MSCs. (k) A cartoon image shows the interaction of DPP4 and GPC3 in N-cad^+^ BMSCs niche. (l) Comparison of the DPP4 enzyme activity of 1 x10^6^ mouse AML cells measured by fluorescence assay with the indicated treatment and time points. Results are reported as relative light units (RLUs) (n = 3 wells). (m) Comparison of CFU capability of DPP4^+/+^ and DPP4^−/−^ AML cells after incubation with or without 100nM GPC3 protein (n = 3 samples). (n) In *in vitro* chemotaxis assays, the migration of AML cells towards CXCL12 induction is significantly reduced in the presence of recombinant GPC3 (100 nM). Error bars, s.e.m. Data were analyzed using a two-tailed Student’s t-test for pairwise analysis. When comparing 3 or more groups, a one-way or two-way ANOVA was used with Fisher’s test for post hoc analyses.

These data suggest that the interaction of GPC3 and DPP4 may mediate the crosstalk between N-cad^+^ MSCs and LSCs (**Fig. 5k**). To investigate the impact of intercellular communication via GPC3 and DPP4 on AML development, GPC3 function was examined. The addition of GPC3 significantly reduced DPP4 enzyme activity, comparable to linagliptin, a DPP4 inhibitor (**Fig. 5l**). Functionally, GPC3 decreased the colony-forming capacity of LSCs in CFU assays, with no effect on *Dpp4^−/−^*AML cells (**Fig 5m**). Moreover, in chemotaxis assays, the migration of *Dpp4^+/+^* AML cells towards CXCL12 was significantly reduced in the presence of recombinant GPC3 (**Fig. 5n**). This reduction in migration mirrored the behavior of *Dpp4^−/−^* AML cells in the same assays (**Extend Data Fig 4e, f**). These findings suggested that GPC3, highly expressed on N-cad^+^ MSCs, serves as an endogenous inhibitor of DPP4, influencing the stemness and trafficking of AML cells.

## Discussion

LSCs present significant treatment challenges, often leading to minimal residual disease and relapse^63^. Prior studies have identified human LSCs predominantly in the endosteal region of trabecular bone (metaphysis) in patient samples and xenograft models ^64–66^, which aligns with our findings of LSC localization in the bone marrow (BM) of control mouse AML models. However, our research reveals distinct localization patterns when examining LSCs deficient in *Dpp4* or in recipients lacking *Cxcl12* in N-cadherin-positive cells, with these LSCs primarily located in the CM area. This divergence in BM microenvironment localizations provides a unique opportunity to explore the dynamic interplay between LSCs and niche cells, offering insights into the mechanisms governing LSC properties.

Traditionally, Cxcl12 has been recognized for mediating HSC and LSC quiescence^67–69^. It was thought that CXCL12 directly induced LSC quiescence, contributing to leukemia relapse. However, our study challenges this notion, revealing that CXCL12 does not directly induce LSC quiescence but rather regulates LSC mobilization between two distinct niches: the metaphysis niche marked by N-cad^+^ MSCs and the central marrow niche marked by sinusoid endothelial cells. We observed that the prior niche where LSCs naturally enrich is crucial for LSC maintenance, with LSCs in the CM region showing proliferation and apoptosis when high levels of CXCL12 are present.

The localization of LSCs to specific niches and their peripheralization is influenced by the concentration of CXCL12, with LSCs tending to migrate toward regions with higher CXCL12 levels. This gradient is regulated by DPP4 and GPC3. LSCs exhibit high expression of DPP4 on their cell membranes, which deactivates CXCL12. Knocking out *Dpp4* results in increased CXCL12 levels in the CM, leading to enriched LSCs in that region. In contrast, the metaphysis region naturally exhibits higher expression of CXCL12 due to N-cad^+^ MSC-specific expression of GPC3, which inhibits DPP4 and reduces CXCL12 deactivation. Thus, targeting DPP4 emerges as a promising strategy to disrupt LSC maintenance.

Notably, DPP4 depletion confines AML cells to the BM, likely due to the overall increased CXCL12 levels in the BM. We observed a reversed gradient of CXCL12 between the BM and PB after *Dpp4* KO, suggesting that higher CXCL12 concentrations in the PB than in the BM are necessary for AML cell peripheralization. This confinement of LSCs to the BM presents a promising therapeutic strategy for AML, potentially reducing tumor burden not only in the PB but also in other organs. This approach offers the prospect of alleviating complications such as leukostasis, clotting abnormalities, respiratory distress, and stroke, thereby reducing associated morbidity ^70,71^. Additionally, targeting the BM niche can activate quiescent LSCs, driving them out of the protective microenvironment and rendering them vulnerable to apoptosis. This reinforces the concept that leukemia cells have a propensity for differentiation, increasing their susceptibility to chemotherapy ^72–75^.

The role of Cxcl12 and its intricate relationship with the BM microenvironment has been a subject of controversy in leukemia development^15,17–19^. While much research has focused on identifying the single cellular niche responsible for *Cxcl12* origin, its relationship with LSC cell proliferation has been less explored. Our study emphasizes the pivotal role of N-cad^+^ MSCs as the primary source of Cxcl12, influencing downstream signaling pathways such as NF-kB, MAPK, and Stat3 in LSCs. Furthermore, our findings elucidate the intricate interaction between DPP4 and GPC3, providing insight into how AML cells interact with the BM niche and suggesting a potential therapeutic target.

In conclusion, our study unravels a complex network involving CXCL12, DPP4, and GPC3 in the BM microenvironment, regulating AML cell trafficking, localization, and microenvironmental-regulated stemness and survival. This novel insight offers a promising avenue for targeted therapeutic interventions aimed at minimal residual LSCs, with the potential to achieve a lasting cure for AML.

## Acknowledgments

We thank the University of Missouri Genomics Core for their support in generating data on the scRNAseq.

## Funding

This work was funded by the National Cancer Institute (Grant R37CA241603 to X.K.), and the American Cancer Society (Grant RSG-23-1152630 to X.K.).

## Author Contributions

X.K. conceived and supervised the study; C.W. performed experiments and analyzed data; Y.P., C.W., W.Z., Y.S., and X.M. conceived the experiments; C.W., W.Z. performed scRNA-seq and bulk RNA-seq. Y.P. performed bioinformatics analysis. C.W. performed immunofluorescence staining and imaging data analysis. C.W., Y.P., W.Z. and X.M. verified the reproducibility of results; R.D.H. contributes to BM imaging data analysis; L.L.,R.D。, R.N. provided technical assistance and contribute to data analysis. Y.P. and X.K. wrote the original draft.

### Declaration of interest

The authors declare no competing interests.

## Notes

### Competing Interest Statement

The authors have declared no competing interest.

